# Female-Specific Pituitary Hypersensitivity to Gonadotropin-Releasing Hormone in a Mouse Model of Chronic Temporal Lobe Epilepsy

**DOI:** 10.1101/2022.11.16.516789

**Authors:** Cathryn A. Cutia, Leanna K. Leverton, Karen E. Weis, Lori T. Raetzman, Catherine A. Christian-Hinman

## Abstract

Gonadotropin hormone release from the anterior pituitary is critical to regulating reproductive endocrine function. Clinical evidence has documented that people with epilepsy display altered levels of gonadotropin hormones, both acutely following seizures and chronically. Despite this relationship, pituitary function remains a largely understudied avenue in preclinical epilepsy research. Recently, we showed that females in the intrahippocampal kainic acid (IHKA) mouse model of temporal lobe epilepsy were found to display changes in pituitary expression of gonadotropin hormone and GnRH receptor genes. Circulating gonadotropin hormone levels, however, have yet to be measured in an animal model of epilepsy. Here, we evaluated the circulating levels of luteinizing hormone (LH) and follicle-stimulating hormone (FSH), GnRH receptor (*Gnrhr*) gene expression, and sensitivity to exogenous GnRH in IHKA males and females. Although no changes in overall dynamics of pulsatile patterns of LH release were found in IHKA mice of either sex, estrus vs. diestrus changes in basal and mean LH levels were larger in IHKA females with prolonged, disrupted estrous cycles. In addition, IHKA females displayed increased pituitary sensitivity to GnRH and higher *Gnrhr* expression. The hypersensitivity to GnRH was observed on diestrus, but not estrus. Chronic seizure severity was not found to be correlated with LH parameters, and FSH levels were unchanged in IHKA mice. These results indicate that although there are changes in pituitary gene expression and sensitivity to GnRH in IHKA females, there may also be compensatory mechanisms that aid in maintaining gonadotropin release in the state of chronic epilepsy in this model.

## Introduction

Epilepsy is a common disorder, affecting approximately 1.2% of the US population (1) and nearly 50 million people worldwide (2). For many people with epilepsy, the diagnosis can be further complicated by the development of conditions comorbid to the recurrent seizures.Reproductive endocrine dysfunction is one such comorbidity, and is seen most commonly in people with temporal lobe epilepsy (TLE), the most prevalent form of focal epilepsy (3). Women with TLE show increased rates of polycystic ovary syndrome (PCOS) and hypogonadotropic hypogonadism compared to the general population (4), whereas men with TLE often show hypogonadotropic hypogonadism (5). Although certain antiseizure medications can elicit endocrine abnormalities (6), substantial clinical evidence highlights that reproductive endocrine dysfunction is not solely attributable to the side effects of these drugs. For example, many women with TLE develop menstrual disorders in the absence of antiseizure medications (4,7), and the location of the seizure focus can impact the occurrence of specific reproductive endocrine comorbidities (4,8,9). This evidence further suggests a relationship between seizure activity and reproductive endocrine dysfunction. Despite the association of TLE and reproductive endocrine dysfunction in people with epilepsy, specific mechanisms linking temporal lobe seizures to reproductive endocrine dysfunction have yet to be elucidated.

Clinical evidence indicates that pituitary gonadotropin release is altered by focal seizures. Levels of both luteinizing hormone (LH) (10) and follicle-stimulating hormone (FSH) (11,12) shift acutely following seizures and chronically in human TLE. This evidence suggests that disruptions in the neuroendocrine circuitry that regulates gonadotropin release are likely candidates for mechanisms linking seizures to comorbid reproductive endocrine dysfunction. LH and FSH are released from the anterior pituitary mainly in response to gonadotropin-releasing hormone (GnRH) input from the hypothalamus (13). The mechanisms of release differ between these two hormones; LH release is pulsatile, whereas FSH release is constitutive (13). In addition, the levels of these hormones relative to one another depend largely on the frequency of GnRH pulsatile input (14). As the pulsatile release of LH largely mimics that of GnRH release (15), characterizing the pulsatile levels of LH in the blood can be indicative of shifts in GnRH input to the pituitary. GnRH signals to pituitary gonadotropes through the GnRH receptor (GnRH-R). The number of these receptors expressed on the cell surface of gonadotropes fluctuates to regulate sensitivity of the pituitary to GnRH at different stages of reproductive development (e.g., before and after puberty), and at different stages of female ovarian cycles, to effectively regulate the amount of gonadotropin hormone released (16,17). Therefore, alterations in gonadotropin release with focal seizures could be due to aberrant GnRH input to the pituitary, changes in pituitary sensitivity to GnRH, or a combination of these two factors.

Several animal models of TLE exhibit signs of reproductive endocrine dysfunction alongside epileptogenesis and chronic epilepsy, particularly in females. For example, female rats in the pilocarpine post-status epilepticus model of TLE often develop PCOS-like symptoms (18), including ovarian cysts and hyperandrogenism. Additionally, multiple rodent models of epilepsy exhibit disrupted estrous cycles (18–22). In the intrahippocampal kainic acid (IHKA) mouse model of TLE, hypothalamic GnRH neurons show disrupted firing and excitability in a manner that not only changes over the estrous cycle, but is also associated with the severity of estrous cycle disruption (23,24). These effects are also reflected in changes in GABAergic transmission to GnRH neurons (25). IHKA females were also recently found to display differences in expression of gonadotropin hormone subunit and GnRH receptor genes in the pituitary (22). These pituitary gene expression changes depended on the side of hippocampus targeted for IHKA injection (22), thus further supporting a role for seizure activity itself in driving these downstream effects. Taken together, these findings indicate that functional shifts in the GnRH-gonadotrope axis are likely mechanisms linking reproductive endocrine dysfunction and seizures, and that these changes can, at least in part, be modeled and studied in IHKA mice.

Although epilepsy-induced changes in circulating gonadotropins may help drive estrous cycle disruption and other forms of reproductive endocrine dysfunction, assessments of gonadotrope function and circulating gonadotropins have yet to be studied in an animal model of epilepsy. Therefore, the present study was conducted to test the hypothesis that pituitary release of LH and FSH, GnRH receptor gene expression, and sensitivity to GnRH input are changed in the IHKA model of TLE in a sex-dependent manner.

## Methods

### Animals

All animal procedures complied with the ARRIVE guidelines and were approved by the Institutional Animal Care and Use Committee of the University of Illinois Urbana-Champaign. GnRH-Cre females and males (Jackson Labs #021207) on the C57BL/6J background were housed in a 14:10 h light:dark cycle (lights off at 1900 h) with food and water available *ad libitum*. The animals used in this study were progeny not used in breeding pairs for ongoing projects that require Cre expression in GnRH neurons (23–25). After weaning, all animals were group-housed with 2-5 mice per cage. Estrous cycle monitoring was performed from 0900 to 1100h as previously described starting at postnatal day (P) 42 (26) (**Figure 1**). Daily cycle stage monitoring was done for at least 2 weeks prior to intrahippocampal injection, and again for 14-17 days prior to initial tail blood sampling. The cycle length was calculated as the time it took on average to progress from one day of estrus through each stage to the next day of estrus (23,27). Cycle lengths of seven days or more were categorized as ‘long’ (23,27).

**Figure 1.**
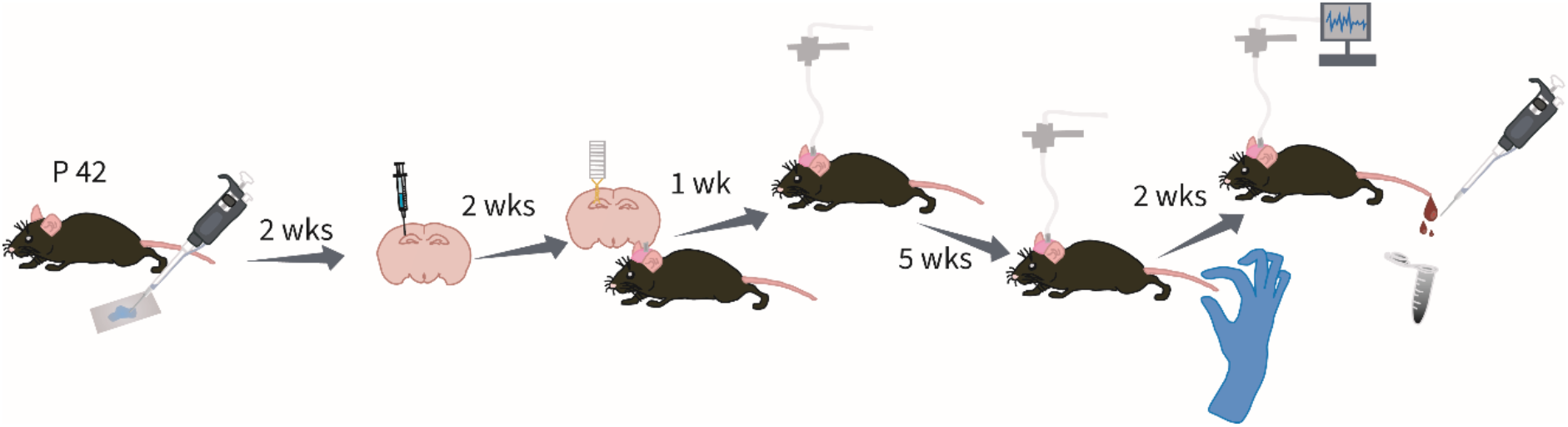
Timeline of experimental procedures. Animals were injected intrahippocampally with KA or saline after 2 weeks of estrous cycle monitoring. Mice were then implanted and habituated to tethering before being habituated to having the tail touched. Mice then had EEG recordings collected and underwent repeated tail blood sampling.

### Intrahippocampal kainic acid injection and EEG electrode implantation

All surgical procedures were conducted under isoflurane inhalation anesthesia. At postnatal day 56 and older, mice underwent unilateral injection of kainic acid (Tocris Bioscience; 50 nl, 20 mM prepared in 0.9% sterile saline) into the right dorsal hippocampal region (from Bregma: 2.0 mm posterior, 1.5 mm lateral, 1.4 mm ventral to the cortical surface). All injections in females were performed on diestrus. Controls were injected with the same volume of saline at the same location. 14 days after injection, the mice were implanted with twisted bipolar EEG electrodes (P1 Technologies) into the ipsilateral hippocampus (22,24,28,29). Implantation was slightly dorsal and lateral to the injection location (from Bregma: 1.75 mm lateral, 1.25 mm ventral). Five anchor micro-screws (J.I. Morris Co.) were then inserted into and distributed across the frontal, parietal, and occipital skull plates. The electrode and screws were stabilized to the skull surface with dental cement (Teets ‘Cold Cure’ Dental Cement). Mice were singly housed after EEG instrumentation to avoid damage to the implant.

### EEG recording and analysis

7 days after EEG electrode implantation, the mice were singly housed and tethered to an electrical commutator (P1 Technologies). The mice were habituated to tethering for five weeks prior to experimentation to avoid tethering-induced disruption to the estrous cycle (29). 24 hours prior to initial tail blood sampling, the commutator was connected to a Brownlee 440 amplifier (NeuroPhase) with the gain set at 1K to collect EEG signals recorded as the local field potential differential between the two electrodes (28). These signals were sampled at 2 kHz and digitized to a computer with recorder software written in MATLAB (28). At 2 months post-injection, the mice were recorded for 9-24 hours prior to tail blood sampling (24) and through the 3-hour sampling period. Recordings were analyzed with an automated electrographic seizure analyzer (30), with the minimal seizure duration set at five seconds and interictal interval set at six seconds.

### Tail blood sampling and trunk blood collection

Beginning 14 days prior to initial tail blood sampling, mice underwent daily tail touch habituation in which the tail was stroked 6 times every 5 minutes for 15 minutes (33). This habituation is necessary to reduce stress, which can affect LH release (31). Mice were sampled twice at approximately 2 months after intrahippocampal injection: once on a day in diestrus and once on a day in estrus. The sampling order of mice on diestrus or estrus was counterbalanced to ensure equal numbers of animals underwent first sampling at each cycle stage. On the day of sampling, <1 mm of the tail tip was clipped off and 6 µl of blood was pipetted off the tail using pipette tips prewashed in EDTA (32) (**Figure 1**). Samples were collected every five minutes for three hours with minimal restraint of the mouse between 0800 h and 1200 h. The blood samples were diluted in 54 µl of buffer (prepared as 0.5 g BSA and 125 µL Tween 20 in 250 mL of 1X PBS) and stored on ice. Mice were returned to their home cage between sample collections. At the end of the 3-hour period of LH sampling, a 150 ng/kg bolus intraperitoneal (IP) injection of GnRH (Bachem 4033013; LHRH acetate salt prepared in 0.9% sterile saline) was administered (‘GnRH challenge’). 15 minutes after this injection, a final tail blood collection was made. Once all samples for a given session were collected, mice were allowed to recover for at least 7 days before further sampling or experimentation. Blood samples were stored at -20°C before being assayed.

Trunk blood, collected at the time of euthanasia via decapitation at least one week after the last blood sampling period, was stored at 4°C overnight, and following centrifugation serum was collected for FSH assays. Sample collection from females was conducted on a day of estrus or diestrus, assigned randomly to each mouse.

### Measurement of blood LH and FSH levels

LH and FSH concentrations were measured by the University of Virginia Ligand Assay and Analysis Core using ultrasensitive enzyme-linked immunosorbent assays (ELISAs) for mouse LH in whole blood (34, 35) for mouse FSH in serum (33). The functional sensitivity of both assays is 0.016 ng/ml.

### Pituitary qPCR

Pituitaries were removed at the time of euthanasia and stored in RNAlater (Invitrogen) at -20°C. RNAqueous-Micro kits (Invitrogen) were used to isolate pituitary RNA. The ProtoScript First Strand cDNA Synthesis kit (New England Biolabs) was then used to reverse transcribe the total RNA (22,34). Oligonucleotide primers for the GnRH receptor (*Gnrhr*) (Life Technologies, **Table 1**) gene were used to amplify gene-specific transcripts by quantitative PCR (qPCR). mRNA expression levels of Peptidylprolyl isomerase A (*Ppia*), a common reference gene used as an internal control, were used to normalize the expression levels of the gene of interest. The data were analyzed using the standard comparative cycle threshold value method (2^−Δ Δ Ct^).

**Table 1.**
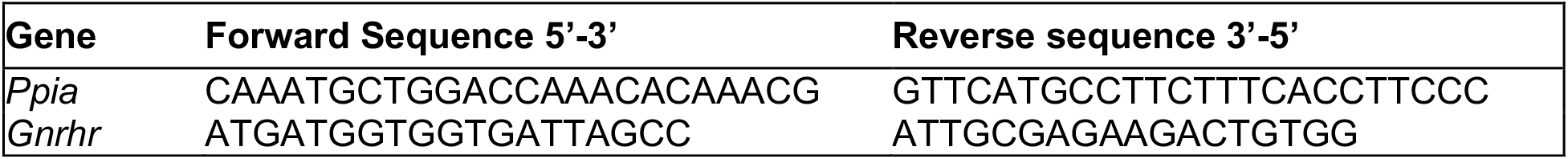
Primer sequences. List of the primer sequences used for qPCR quantification of *Gnrhr* and control *Ppia*.

### Evaluation of LH pulse parameters

LH peaks and pulse dynamics were quantified in a blinded manner. LH concentration values that showed at least a 20% increase from prior points and were followed by an immediate decrease of at least 10% were characterized as pulse peaks as previously described (31,35). All other parameters (amplitude, basal levels, and mean release) were calculated for each mouse using established standards (31,35). Specifically, LH pulse amplitude was calculated as the difference between the concentration of an LH peak, and the trough (lowest value) immediately preceding the peak. Basal LH was defined as the average of all LH samples not identified as peak points of release, and mean LH was defined as the average LH concentration across all samples. Interpulse interval was defined as the time between two successive peaks.

### Statistical analyses

Basal, mean, and pulse amplitude measures of LH, post-GnRH challenge differences, and FSH levels were compared between IHKA-long, IHKA-regular, and saline-injected control mice on both diestrus and estrus using one-way ANOVAs. Tukey’s *post hoc* tests were used for further pairwise comparisons. Interpulse interval values were compared between saline/IHKA females on diestrus and estrus using Kruskal-Wallis tests. All comparisons of LH pulse parameters, post-GnRH challenge differences, and FSH levels in males were conducted using two-sample t-tests. Comparisons of percent change between diestrus and estrus for LH pulse parameters and post-GnRH challenge differences were conducted using Kruskal-Wallis tests. Diestrus-to-estrus repeated measures comparisons of LH parameters and post-GnRH challenge differences within each group of females were made using paired t-tests. Comparisons between saline/IHKA males and diestrous females for LH pulse parameters, post-GnRH challenge differences, and FSH levels were made using two-way ANOVAs and Tukey’s *post hoc* tests.

Comparison of *Gnrhr* mRNA expression between saline and IHKA females was made using a Wilcoxon rank sum test. Normality was evaluated with Shapiro-Wilks tests and homogeneity of variance was assessed with Levene’s tests. Statistical significance was set at p < 0.05.

## Results

### Pulsatile release of LH is unchanged in IHKA mice

To evaluate the pulsatile LH release patterns at different stages of the estrous cycle, repeated tail blood samples were collected over two three-hour periods, one when the mice were in diestrus and one during estrus (**Figure 2A-B**). LH concentrations were evaluated for pulsatile dynamics in each treatment group (control n = 10, IHKA-regular n = 6, IHKA-long n = 9 mice). Diestrous females showed no differences in basal LH levels (F(2,22)= 1.59, p = 0.22), mean LH levels (F = 0.91, p = 0.42), LH pulse amplitude (F = 1.05, p = 0.37), or interpulse interval (IPI) (p = 0.29, Kruskal-Wallis) (**Figure 2C**). Similar patterns were seen on estrus, as there were no differences in basal LH levels (F(2,22) = 2.32, p = 0.12), mean LH levels (F = 3.02, p = 0.07), LH pulse amplitude (F = 2.22, p = 0.133), or IPI (p = 0.14, Kruskal-Wallis) (**Figure 2D**) between control, IHKA-long, and IHKA-regular groups.

**Figure 2.**
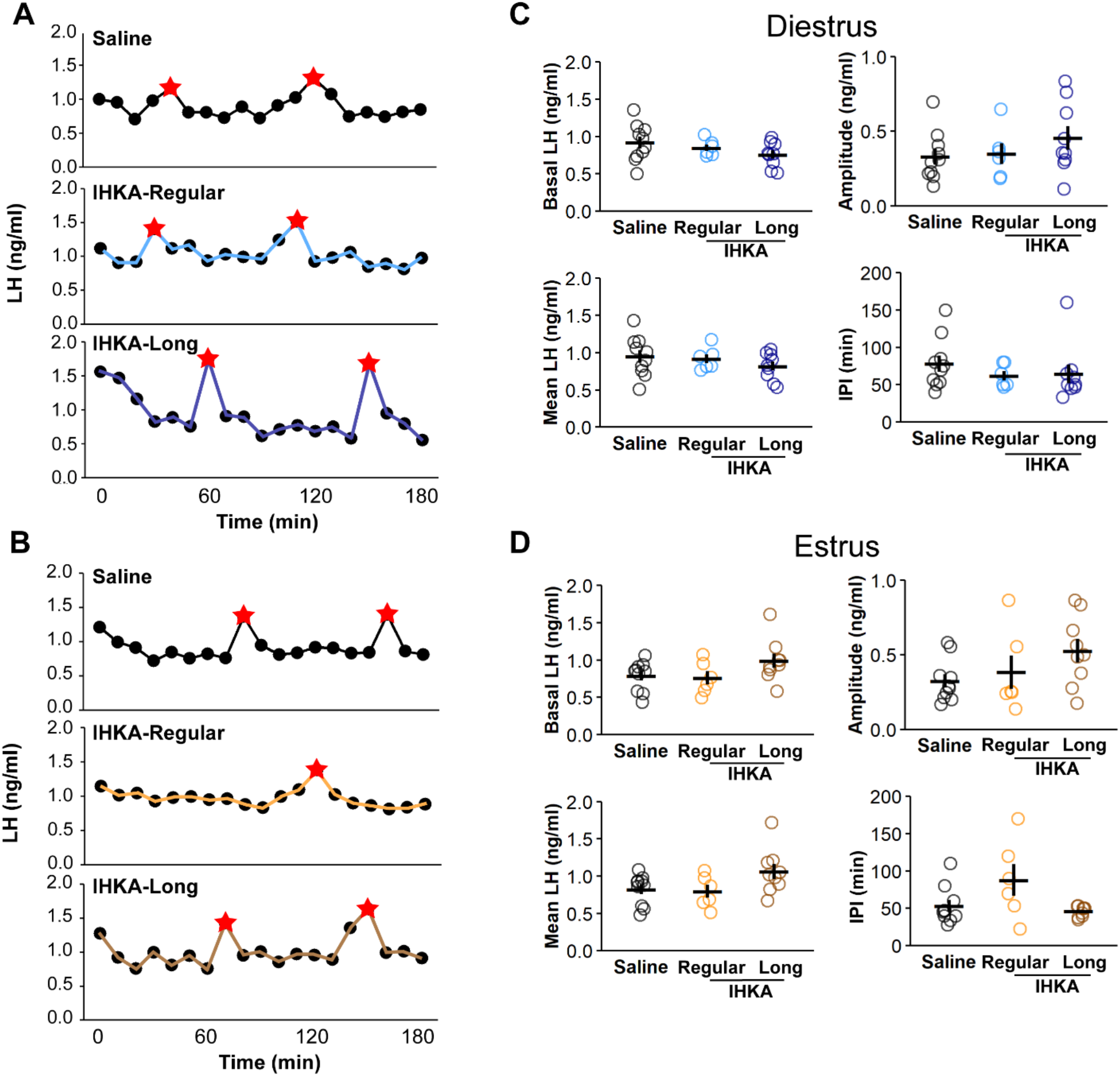
Pulsatile LH release is unchanged in IHKA females. Representative LH pulses from diestrous (A) and estrous (B) animals. Individual values and means ± SEM of mean LH (top left), basal LH (bottom left), pulse amplitude (top right), and interpulse interval (IPI) (bottom right) on diestrus (C) and estrus (D).

To understand if pulsatile dynamics shift in male mice in response to IHKA injection, 11 males (control n= 5, IHKA n = 6) were evaluated (**Figure 3A**). In comparison to controls, IHKA males showed no differences in basal LH levels (t = -0.06, p = 0.95, **Figure 3B**), mean LH levels (t = -0.3, p = 0.77, **Figure 3C**), LH pulse amplitude (t = -0.318, p = 0.76, **Figure 3D**), or IPI (t = 0.66, p = 0.53, **Figure 3E**). The LH values from these males were also compared to diestrous females to test for sex differences in LH pulse parameters. Data from IHKA-long and -regular females were collapsed together for this analysis. While there were no significant differences based on sex in basal LH levels (F(1,32) = 3.66, p = 0.07, **Figure 4A**) or IPI (F = 0.935, p = 0.341, **Figure 4D**), there were significant effects of sex on mean LH levels (F = 6.75, p = 0.01, **Figure 4B**) and LH pulse amplitude (F = 24.62, p = 2.22 × 10^−5^, **Figure 4C**). Specifically, IHKA males showed increased mean LH compared to IHKA females (p = 0.007, **Figure 4B**), and higher LH pulse amplitude compared to both IHKA (p = 0.02) and control females (p = 0.005, **Figure 4C**). LH pulse amplitude was also higher in control males compared to both IHKA (p = 0.008) and control females (p = 0.002, **Figure 4C**).

**Figure 3.**
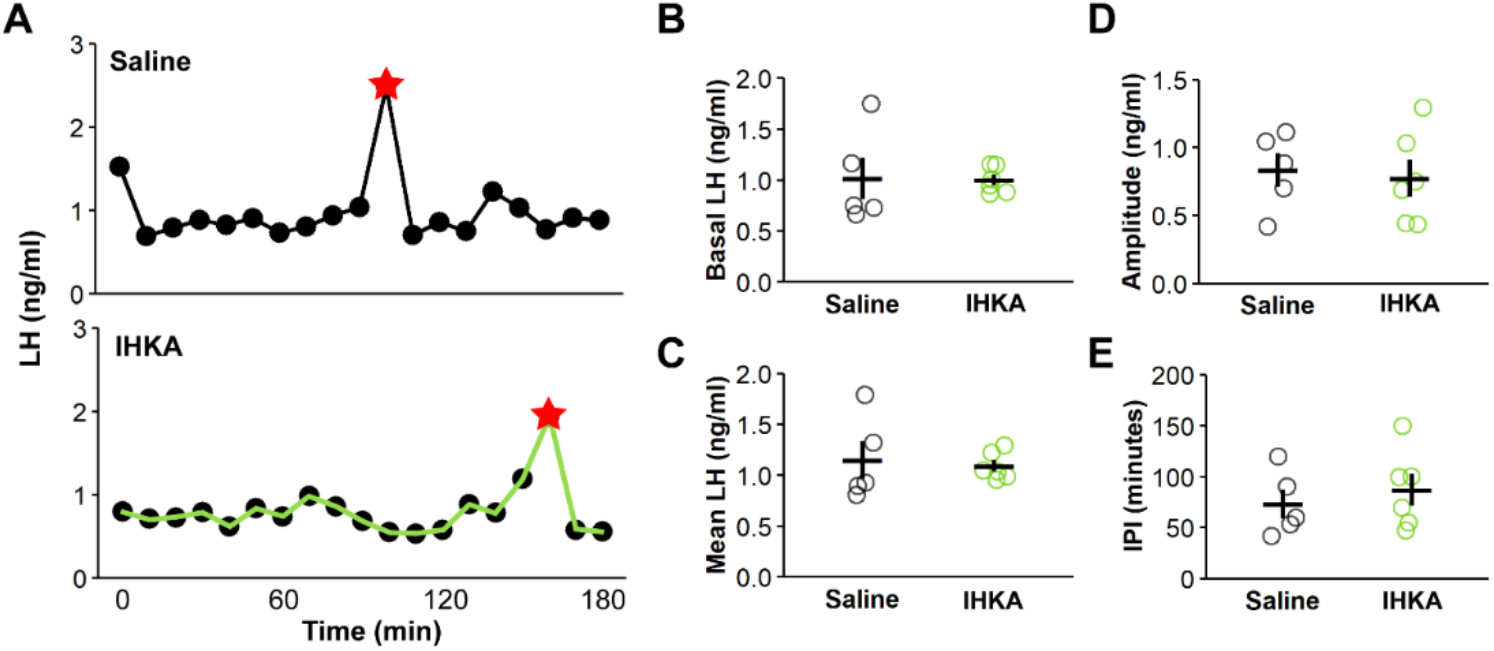
Pulsatile LH release is unchanged in IHKA males. (A) Representative LH traces from saline (top) and IHKA (bottom) males. Individual values and means ± SEM of basal LH (B), mean LH (C), pulse amplitude (D), and interpulse interval (IPI) (E).

**Figure 4.**
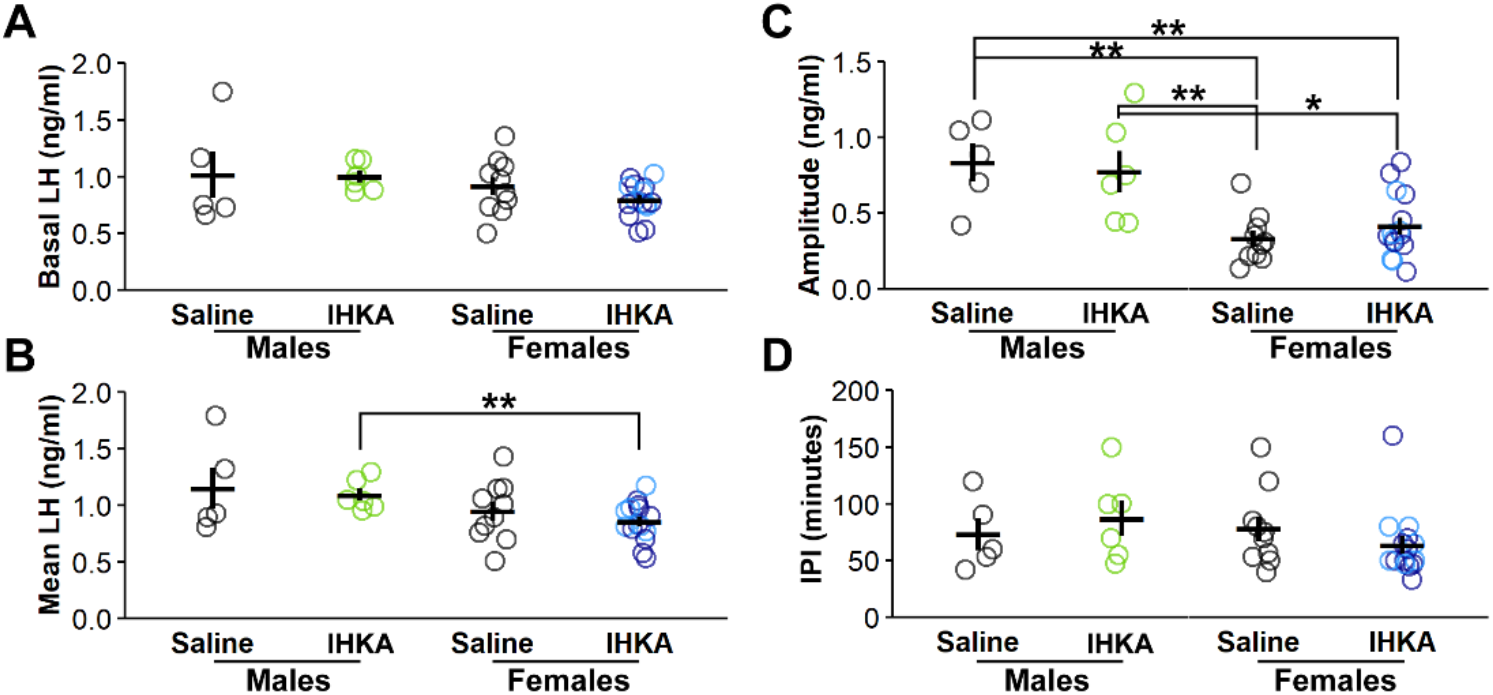
LH pulse amplitude is larger in males compared with females. Individual values and means ± SEM of mean LH (A), basal LH (B), pulse amplitude (C), and interpulse interval (IPI) (D). ***p < 0.001 for comparisons between sex (male and female) and treatment (IHKA and saline) by two-way ANOVA and Tukey’s *post hoc* tests.

### Increased estrus vs. diestrus changes in mean and basal LH levels in IHKA-long mice, but not IHKA-regular or control groups

To evaluate whether there were significant shifts in LH release between diestrus and estrus, and whether these cycle-associated changes differ between control and IHKA females, data collected from the same mice on diestrus and estrus were evaluated for each LH parameter. Comparisons within saline controls showed that basal LH (t = 1.63, p = 0.13, **Figure 5A**), mean LH (t = 1.46, p = 0.18, **Figure 5B**), LH pulse amplitude (t = 0.11, p = 0.92, **Figure 5C**), and IPI (t = 1.66, p = 0.13, **Figure 5D**) were not different on estrus compared with diestrus. IHKA-regular animals also showed no significant changes from diestrus to estrus in terms of basal LH (t = 1.84, p = 0.1253, **Figure 5A**), LH pulse amplitude (t = -0.38, p = 0.72, **Figure 5C**), or IPI (t = -1.21, p = 0.28, **Figure 5D**), but mean LH levels were reduced in IHKA-regular mice on estrus compared with diestrus (t = 3.49, p-value = 0.02, **Figure 5B**). IHKA-long mice, similarly to controls, showed no significant changes from diestrus to estrus in terms of basal LH (t = -2.09, p = 0.07, **Figure 5A**), mean LH (t = -2.01, p = 0.08, **Figure 5B**), LH pulse amplitude (t = -0.59, p = 0.57, **Figure 5C**), or IPI (t = 1.51, p = 0.17, **Figure 5D**). However, comparisons of the percent change in basal and mean LH levels from diestrus to estrus showed that there were differences between the groups (basal LH p = 0.008; mean LH p = 0.02, Kruskal-Wallis, **Figure 5A, 5B**), as IHKA-long mice showed larger estrus vs. diestrus changes compared to both controls (basal LH p = 0.009; mean LH p = 0.03, Dunn’s, **Figure 5A, 5B**) and IHKA-regular mice (mean LH p = 0.04, Dunn’s, **Figure 5B**). LH pulse amplitude (p = 0.82, **Figure 5C**) and IPI (p = 0.51, **Figure 5D**) did not show any cycle stage-dependent shifts.

**Figure 5.**
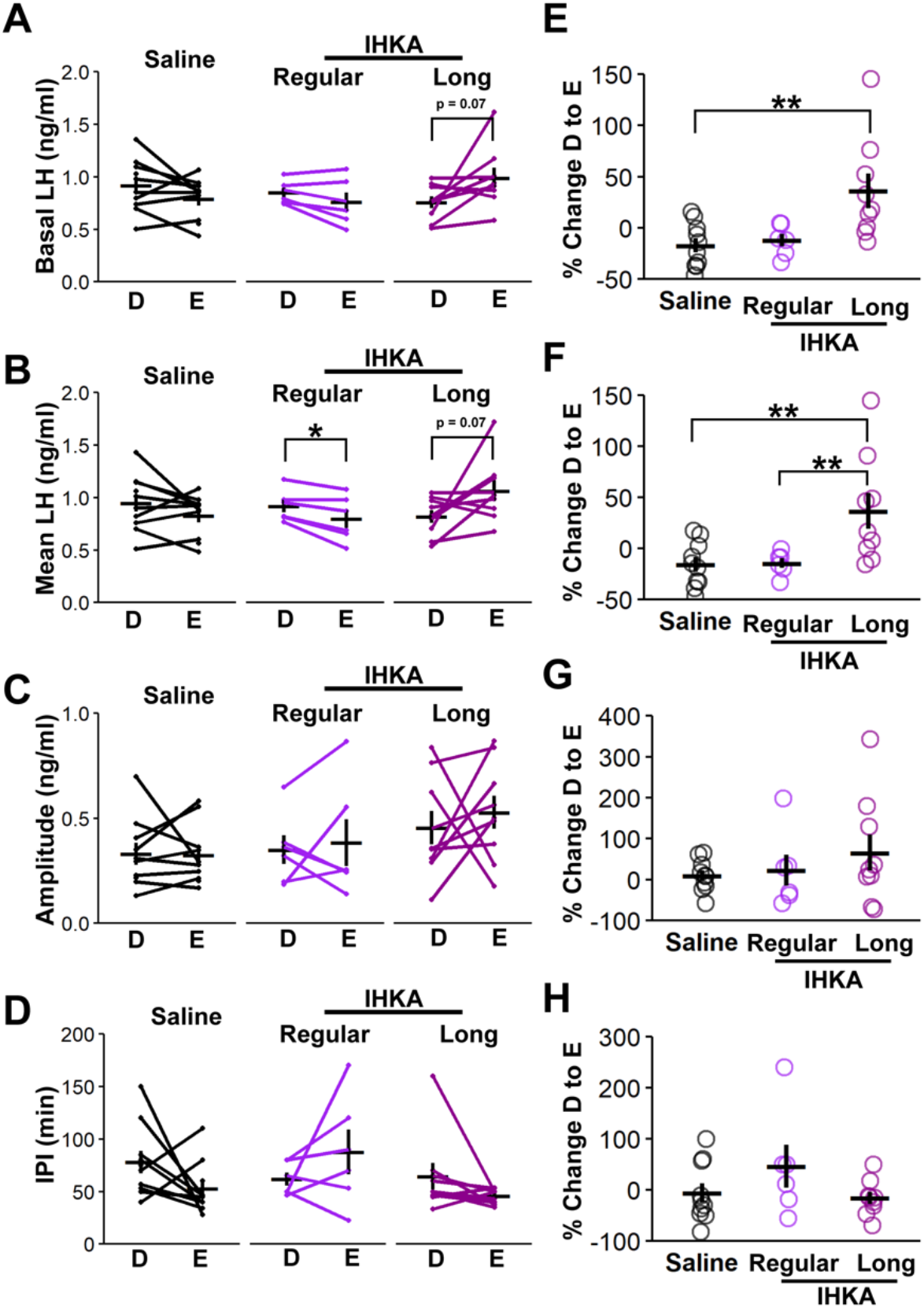
Basal and mean LH levels increase in IHKA-long animals from diestrus to estrus. A-D) Representative changes in basal LH (A), mean LH (B), LH pulse amplitude (C), and IPI (D) plotted with mean ± SEM. *p<0.05 for comparisons of diestrus to estrus by a paired t-test. E-H) Individual values of the percent change from diestrus to estrus in basal LH (E), mean LH (F), LH pulse amplitude (G), and IPI plotted with mean ± SEM. **p<0.01 Kruskal-Wallis and Dunn’s tests

### Seizure burden is not correlated to changes in LH release

Previous studies have shown that 24 hours of EEG monitoring is an accurate approximation of overall seizure burden in IHKA mice (24). Therefore, to confirm the development of chronic epilepsy and to evaluate whether seizure burden in the 24-hour period prior to LH collection was correlated to pulsatile LH release patterns, we compared EEG signals to the respective LH parameters from each animal. Control mice showed no seizures, whereas all IHKA mice showed at least two electrographic seizures during this recording period. There were no correlations between percent time spent in seizures and LH pulse amplitude, mean LH levels, or the IPI in either estrous or diestrous females, or in males (**Table 2**).

**Table 2.** LH pulse parameters do not correlate to seizure burden. r^2^ and p values for Pearson correlations between mean LH levels, LH pulse amplitude, and interpulse interval made to the percent time spent in seizures quantified in 24 hours prior to tail blood sampling.

|  | % time spent in seizures |  |  |  |  |  |
| --- | --- | --- | --- | --- | --- | --- |
|  | *Diestrus* |  | *Estrus* |  | *Males* |  |
|  | *r*<sup>2</sup> | *p* | *r*<sup>2</sup> | *p* | *r*<sup>2</sup> | *p* |
| **Mean LH levels** | -0.24 | 0.37 | -0.17 | 0.52 | -0.48 | 0.34 |
| **LH pulse amplitude** | -0.08 | 0.77 | 0.21 | 0.43 | -0.06 | 0.91 |
| **Interpulse interval** | -0.29 | 0.28 | 0.10 | 0.71 | -0.29 | 0.58 |

### Increased sensitivity to exogenous GnRH in diestrous IHKA females

Under normal conditions, LH is released from the pituitary in response to GnRH input. To test whether pituitary sensitivity to GnRH is changed in animals with epilepsy, tail blood was collected 15 min after administration of a bolus IP injection of GnRH (‘GnRH challenge’) (36,37) at the end of each baseline 3-h sampling period (**Figure 6A**). The difference in LH concentrations from tail blood samples collected prior to and 15 minutes following the GnRH injection was calculated for each mouse (**Figure 6B**) and compared between groups (**Figure 6C-E**). Diestrous females showed a significant change in this difference based on IHKA vs. control treatment (F(2, 22) = 5.02, p = 0.02, **Figure 6D**); specifically, *post hoc* analysis revealed that IHKA-long females exhibited a larger pre-to post-injection increment in LH levels compared to controls. IHKA-regular animals, however, did not show the same effect in comparison to controls (p = 0.71, **Figure 6D**). By contrast, post-GnRH LH increment was not different in estrous IHKA females compared to corresponding controls (F=0.841, p = 0.45, **Figure 6E**). Similarly, control and IHKA males displayed no differences in post-GnRH LH increment (t = - 0.83, p = 0.43, **Figure 6C**).

**Figure 6.**
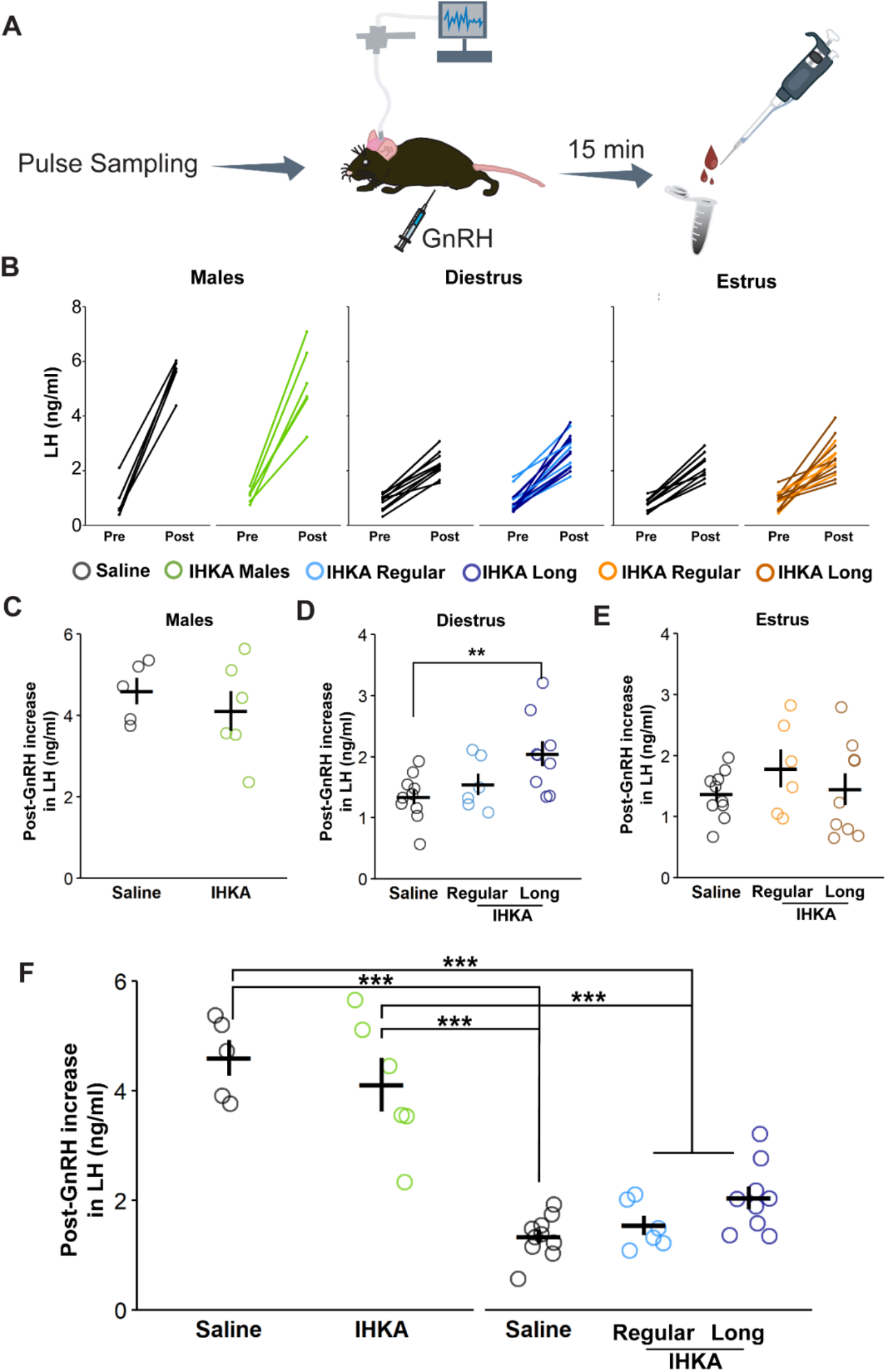
Female-specific increases in pituitary sensitivity to exogenous GnRH. (A)Timeline of experimental methods for evaluating GnRH sensitivity. (B) LH values for each animal prior to and following IP injection of GnRH. (C-E) Individual values in LH differences following GnRH injection in males (C), diestrous females (D), and estrous females (E). (F) Comparisons of individual values from males and diestrous females. All individual values are presented with mean ± SEM values. ***p < 0.001 for comparisons between sex (male and female) and treatment (IHKA and saline) by two-way ANOVA and Tukey’s *post hoc* tests.

To determine whether pituitary sensitivity to GnRH in control and/or IHKA mice exhibits sex differences, post-GnRH LH increment values in males were compared to values from diestrous females. The post-injection LH increment was significantly higher in males compared to females (F(1,32) = 113.31, p = 4.81 × 10^−12^) regardless of treatment type (**Figure 6F**). Specifically, IHKA males showed larger post-GnRH LH increments compared to both IHKA (p = 8 × 10^−7^) and control females (p = 1 × 10^−7^), and control males showed higher values in comparison to both IHKA (p = 1 × 10^−7^) and control females (p = 1 × 10^−8^).

### Estrous cycle stage does not impact LH release after systemic GnRH injection

To determine if estrous cycle stage impacts the pituitary response to the GnRH challenge, the difference in LH from pre-to post-injection on diestrus was compared to estrus using paired statistics for each group. Controls showed no change in the mean response on diestrus compared to estrus (t = -0.22, p = 0.83, **Figure 7A**). Similarly, IHKA-regular mice showed no significant change with cycle stage (t= -0.52, p = 0.63, **Figure 7A**). IHKA-long mice, however, showed a trend for a decrease in this response from diestrus to estrus (t = 1.98, p = 0.08, **Figure 7A**). Comparisons were then made between the percent change in these values on diestrus to estrus for each of the groups. Although the clear majority of IHKA-long mice showed a reduction in response on estrus compared with diestrus, there were no differences between control and IHKA females (p = 0.13, Kruskal-Wallis, **Figure 7B**).

**Figure 7.**
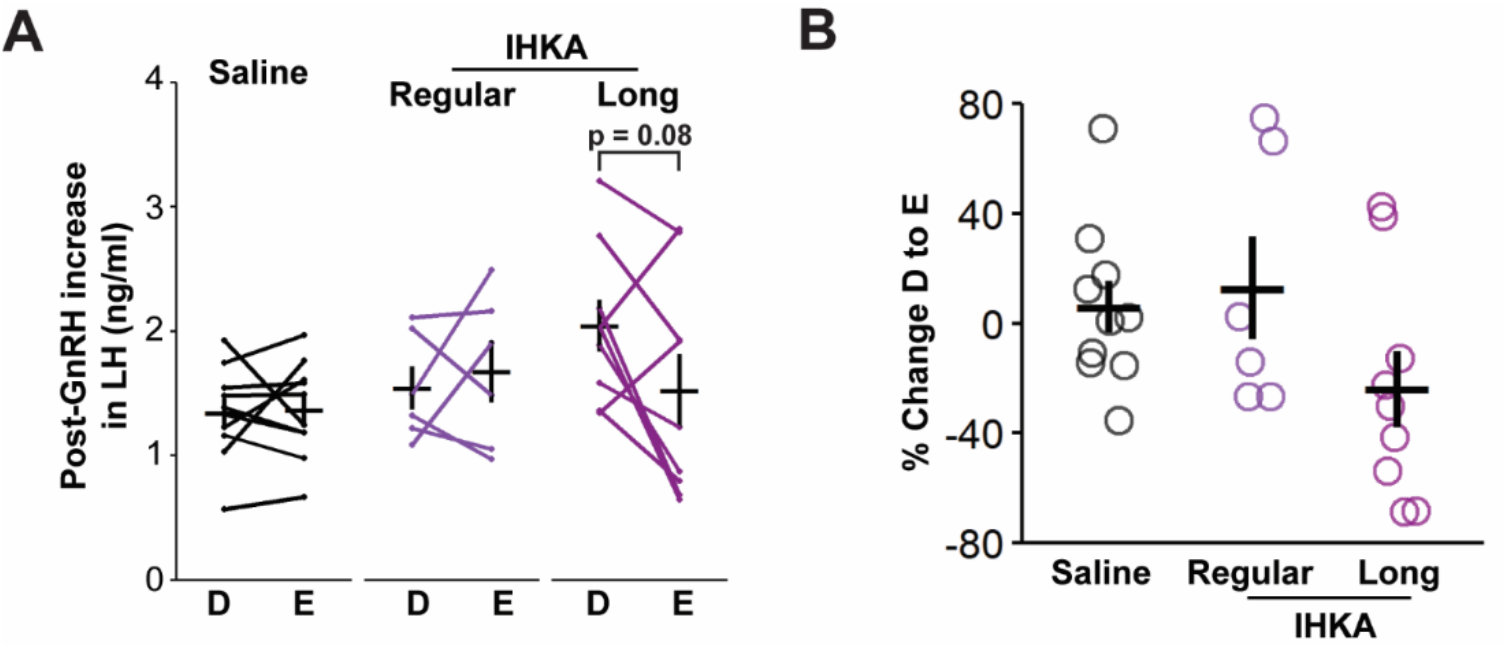
IHKA-long mice, but not controls nor IHKA-regular mice, display trend towards decreased pituitary sensitivity to GnRH on estrus compared with diestrus. (A) Representative changes in LH following GnRH injection for each animal on diestrus and estrus. (B) Individual values of the percent change in post-GnRH LH increment from diestrus to estrus plotted with mean ± SEM values.

### Pituitary gene expression of Gnrhr is elevated in female IHKA mice

To understand whether altered GnRH receptor expression might underlie the female-specific increase in sensitivity to GnRH after IHKA injection, mRNA expression levels of the GnRH receptor gene (*Gnrhr*) were evaluated in a subset of female mice for which the collected pituitaries were of suitable quality (saline n = 4, IHKA n = 6). Pituitaries collected on diestrus and estrus were combined due to limited sample size (saline diestrus = 2; saline estrus = 2; IHKA diestrus = 3; IHKA estrus = 3). IHKA mice showed higher *Gnrhr* expression compared to controls (p = 0.01, **Figure 8**), suggesting that pituitary sensitivity to GnRH may be increased through upregulation of GnRH receptor expression in IHKA females.

**Figure 8.**
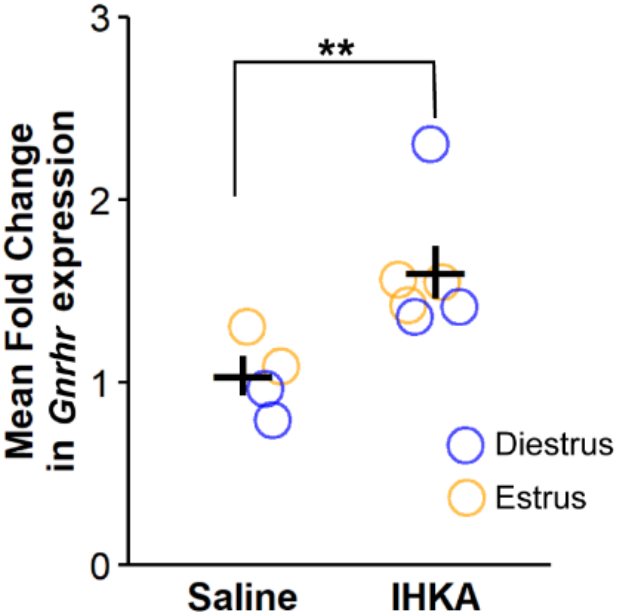
Pituitary *Gnrhr* mRNA expression is increased in IHKA females. Individual values of mean fold change in *Gnrhr* plotted with mean ± SEM. Values from pituitaries collected on diestrus are plotted in blue, while those collected on estrus are plotted in orange. **p≤ 0.01 for comparison between injection type by Wilcoxon rank test.

### FSH levels are unchanged in IHKA mice

Although pulsatile patterns of LH release are more directly associated with GnRH pulses, FSH levels are also impacted by GnRH input to the anterior pituitary. To measure basal levels of FSH in IHKA mice and controls, trunk blood serum from male (saline = 5, IHKA = 6) and female (saline = 9, IHKA-regular = 6, IHKA-long = 7) mice was collected and assayed. Samples from females were collected on diestrus (saline n = 4, IHKA regular n = 3, IHKA long n = 3) or estrus (saline n = 5, IHKA regular n = 3, IHKA long n = 4). Values from diestrous females showed no differences in FSH levels between groups (F(2,7) = 1.33, p = 0.32, **Figure 9A**). Similarly, estrous females (F(2,9) = 0.59, p = 0.58, **Figure 9B**) showed no differences based on cycle elongation phenotype nor injection type, and males (t = 0.32, p = 0.75, **Figure 9C**) showed no differences based on injection type. However, FSH levels were higher in males compared to diestrous females (F(1,17) = 114.77, p = 5.59 × 10^−9^), and in IHKA males compared to both IHKA (p = 1 × 10^−6^) and control (8.4 × 10^−6^) females (**Figure 9D**). FSH was also higher in control males compared to both IHKA (p = 3.4 × 10^−6^) and control (2.4 × 10^−5^) females (**Figure 9D**).

**Figure 9.**
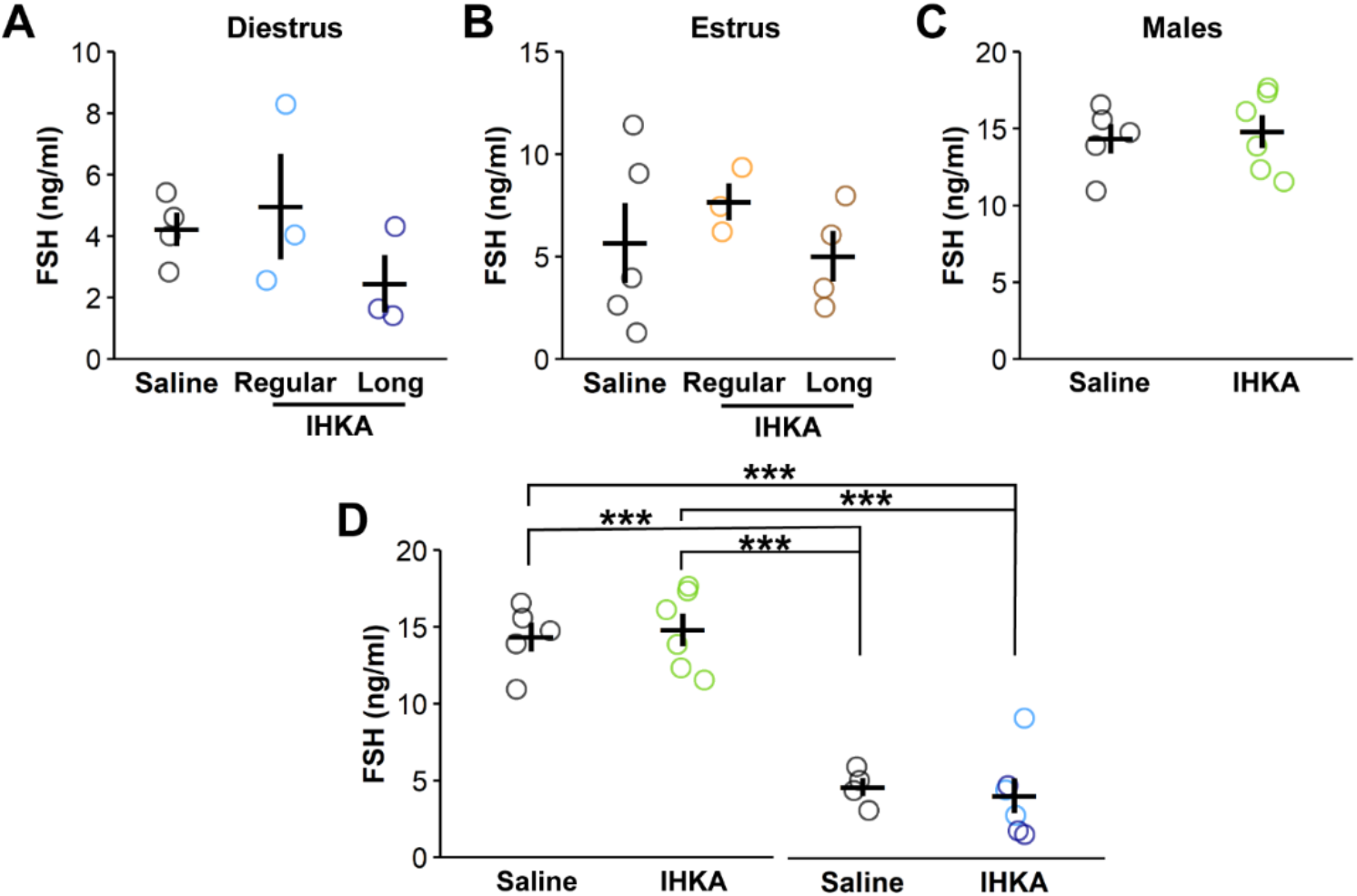
FSH levels are unchanged in IHKA mice. (A-C)Individual values and means ± SEM of serum FSH concentrations for diestrous females (A), estrous females (B), and males (C). (D) Comparisons of FSH values between males and diestrous females. ***p < 0.001 by two-way ANOVA and Tukey’s *post hoc* tests.

## Discussion

Clinical evidence suggests that chronic and acute seizures impact gonadotropin levels and release patterns. Despite this relationship, changes in gonadotropin levels have yet to be documented in a preclinical model of chronic epilepsy. Therefore, in the present study we evaluated LH pulse dynamics, GnRH sensitivity, *Gnrhr* pituitary expression, and basal FSH levels in the IHKA mouse model of TLE. The results indicate that gonadotropin levels are largely unchanged in IHKA mice. However, IHKA females with elongated estrous cycles show a more pronounced estrus vs. diestrus difference in mean and basal LH levels and exhibit higher sensitivity to exogenous GnRH on diestrus. Notably, this effect was not observed in IHKA females that maintained regular cycles. Additionally, we recapitulate previous findings that there is a basal sex difference in the levels of circulating gonadotropins in mice. This result suggests that IHKA female mice with chronic epilepsy and comorbid reproductive endocrine dysfunction may have changes in the GnRH-pituitary axis leading to both altered pituitary sensitivity and compensation to maintain normal gonadotropin hormone release.

Gonadotropin hormones impact the release of gonadal hormones (13). Although gonadotropin levels have not previously been evaluated in a preclinical epilepsy model, gonadal hormone levels have been measured in several models of epilepsy (20). With respect to the IHKA mouse model, the results have been conflicting. In one study using C57BL/6J mice, the background of the GnRH-Cre strain used in the present studies, IHKA females were not found to exhibit changes in estradiol, progesterone, or testosterone when evaluated on diestrus (22). By contrast, another study using mice produced from a cross of GnRH-Cre and the Ai9 Cre-dependent tdTomato reporter strain demonstrated decreased progesterone levels on diestrus and estrus in IHKA-long mice, and elevated estradiol on estrus in both IHKA-long and -regular groups (23). The present results indicate no changes in overall LH or FSH, which would suggest that gonadal hormone levels would, in turn, not be altered. However, the elevated change in basal and LH levels from diestrus to estrus within IHKA-long animals could be indicative of changes in normal patterning of LH release across the cycle, which could contribute to altered downstream function in animals with epilepsy. Additionally, the possibility of changes in ovarian sensitivity to gonadotropin signaling cannot be excluded.

Although there were no differences observed in LH pulsatility or FSH levels, most IHKA females develop elongated estrous cycles (22–24,27). The present results suggest that altered gonadotropin levels may not contribute to the development of this phenotype. Of note, other mouse models, including a model of chronic stress (38) and one of chronic androgen excess (39), also exhibit altered estrous cycles despite a lack of change in LH pulses. Furthermore, LH is released in pulses in both males and females, but once in every ovarian cycle, a surge of GnRH/LH release provides the trigger for ovulation (40). This mechanism of release, known as the preovulatory surge, has similar yet distinct mechanisms compared to LH pulsatile release, and has yet to be measured in IHKA females. It is possible that compensation that occurs to maintain gonadotropin release at more ‘basal’ phases of the estrous cycle may not function as well at the time of the preovulatory surge.

Other studies investigating LH pulsatile release over the estrous cycle have noted changes in pulse parameters with estrous cycle stage (41,42), and demonstrated decreased LH release on estrus compared to diestrus. To some extent, this effect is recapitulated in the decreased mean LH levels on estrus compared with diestrus observed in the IHKA-regular group, although this pattern was not recapitulated in the corresponding saline-injected controls. Of note, the mice in the previous studies were kisspeptin-Cre transgenic mice, whereas in the present study, GnRH-Cre mice were used. In addition, the animals in the present study were housed on a 14:10 light-dark cycle rather than a 12:12 light-dark cycle, and sampling in the present study was carried out over a slightly wider range of time (0800 h to 1200 h) compared to the previous studies, conducted between 0900 h and 1100h. These methodological and strain differences may help, at least in part, to explain these discrepancies.

All IHKA mice included in the present study were confirmed in EEG to have chronic seizures, but no correlations between seizure parameters and LH dynamics were observed. LH release, in particular, is known to be affected by stress in rodents (43), and it is thus possible that the stress of sampling and EEG tethering could impact LH release. All mice in the present study, however, were habituated to the tail sampling process as a means to limit stress-induced suppression of LH (44), and to EEG tethering for over 5 weeks, a duration of time that is sufficient to allow for restoration of estrous cycle regularity after transient tethering-induced suppression (24). It is thus unlikely that EEG tethering and recording influenced the LH measurements here. Although aging influences LH levels and release patterns in mice (45), the mice in the present study were examined at 4 to 5 months of age.

Higher pituitary sensitivity to GnRH in IHKA females in the absence of changes to LH pulses, as highlighted by the elevated responses to exogenous GnRH on diestrus, suggests that the pituitary may compensate for aberrant GnRH input as a way to maintain normal circulating levels of gonadotropins, and thus sustain reproductive endocrine function, in the presence of epilepsy. The higher *Gnrhr* mRNA expression in diestrous and estrous IHKA females supports and extends previous work in diestrous mice only (22), and indicates that gonadotrope expression of GnRH receptors is elevated in IHKA females. This effect could be due to changes in GnRH input to the anterior pituitary (46), or reflect changes in gonadal hormone feedback onto the pituitary (47–49). Due to the small size of the pituitary and the limits of sample numbers, GnRH-R protein was not assessed in the present study. However, the number of membrane GnRH-Rs is, at least in part, dependent on the level of *Gnrhr* mRNA (50). Therefore, it appears reasonable to speculate that increases in the number of GnRH receptors on the gonadotrope surface may underlie the increased sensitivity of the pituitary to GnRH input in IHKA mice. Overall, this result suggests that signals received by the pituitary alter the transcription of the *Gnrhr* gene in IHKA females and may contribute to higher levels of GnRH-R, and thus increase pituitary sensitivity to GnRH.

As noted above, the pituitary contains mechanisms by which membrane expression of GnRH-R, and thus sensitivity to GnRH, shifts based on changing input. For example, GnRH-R levels fluctuate to modulate pituitary sensitivity to GnRH across the rodent estrous cycle (51), and estrogens augment GnRH-R expression to enhance pituitary sensitivity to GnRH during the preovulatory GnRH/LH surge in rodents (48,49). However, GnRH signaling alone can produce increases in *Gnrhr* mRNA (47), although downregulation of GnRH-R expression with chronic exposure to GnRH analogs demonstrates the propensity of the pituitary to become desensitized to GnRH (52,53). Previous studies have shown that GnRH neuron firing is elevated in diestrous IHKA-long females (23). While elevated GnRH neuron firing may result in elevated GnRH release, the aforementioned neural recordings were conducted at the soma of the GnRH neurons, and there is evidence that GnRH neurons differentially receive and integrate inputs at both the soma and the terminals to ultimately lead to GnRH release (54). Additionally, GnRH release in the preoptic area, known to contain GnRH neuron somata, is distinct compared to release at the median eminence, the location of the vast majority of GnRH neuron terminals (55). Therefore, it is possible that firing at the level of the soma does not directly recapitulate the amount of GnRH that is released and interacting with the anterior pituitary. Furthermore, although LH is often used as a readout of GnRH release due to the difficulty of directly measuring GnRH at the median eminence, particularly in small rodents, LH does not always directly reflect GnRH release, particularly during periods of high frequency GnRH release (56,57). Taken together, these findings suggest that there is disruption in the hypothalamic GnRH neuron circuitry in the IHKA model, and this disruption may ultimately result in changed GnRH release, triggering compensated pituitary function as a means of maintaining typical gonadotropin release.

Overall, these results demonstrate that IHKA female mice that develop elongated estrous cycles display elevated sensitivity to GnRH on diestrus and altered dynamics of cycle-associated changes in mean and basal LH levels. This phenotype occurs in the absence of changes to LH pulsatile dynamics. This result, in tandem with previous reports of disruptions in the function of hypothalamic GnRH neurons in IHKA-long mice on diestrus, suggests that there are disruptions occurring in the hypothalamic-pituitary axis due to chronic seizures that are sex- and estrous cycle stage-dependent. Further work is needed to specifically determine how hypothalamic release of GnRH itself is altered, and how epilepsy-associated changes in this axis may contribute to the disrupted estrous cycles observed in this model of epilepsy.

## Acknowledgements

We thank Jiang Li and Jordyn Robare for performing pilot studies establishing the tail blood sampling procedures in our laboratory.

